# Strategies of grain number determination differentiate barley row-types

**DOI:** 10.1101/2020.08.31.275446

**Authors:** Venkatasubbu Thirulogachandar, Ravi Koppolu, Thorsten Schnurbusch

## Abstract

Gaining knowledge on intrinsic interactions of various yield components is crucial to improve the yield potential in small grain cereals. It is well known in barley that increasing the grain number (GN) preponderantly improves their yield potential; however, the yield components determining GN and their association in barley row-types are less explored. In this study, we assessed different yield components like potential spikelet number (PSN), spikelet survival (SSL), spikelet number (SN), grain set (GS), and grain survival (GSL), as well as their interactions with GN by using a selected panel of two- and six-rowed barley types. Also, to analyze the stability of these interactions, we performed the study in two growth conditions, greenhouse and field. From this study, we found that in two-rowed, GN determination is strongly influenced by PSN rather than SSL and/or GS in both growth conditions. Conversely, in six-rowed, GN is associated with SSL instead of PSN and/or GS. Thus, our study exemplified that increasing GN might be possible by augmenting PSN in two-rowed genotypes, while for six-rowed genotypes, the ability of SSL needs to be improved. We speculate that this disparity of GN determination in barley row-types might be due to the fertility of lateral spikelets. Collectively, this study revealed that the GN of two-rowed largely depends on the developmental trait, PSN, while in six-rowed, it mainly follows the ability of SSL.

**Highlight:** In cereals, understanding the interactions of different yield components that influence the grain number is essential to increase their yield by modulating the components. We show in this study that the grain number of two-rowed barley is predominantly determined by the potential spikelet number while in six-rowed by spikelet survival.

## Introduction

Propelling the crop yield is one of the essential aims in the field of plant sciences with the limitations of climatic change and declining arable land. However, crop yield is a complex trait influenced by many abiotic and biotic factors, as well as different plant developmental traits. Barley (*Hordeum vulgare* L.) is the fourth economically important cereal crop in the world (FAOSTAT, 2019). It stands as a model crop of the tribe Triticeae, which contains other major cereals: wheat (*Triticum* spp.), rye (*Secale cereale* L.), and triticale (x *Triticosecale* Wittm.) (Ullrich, 2011; Mascher et al., 2017). Triticeae species have a characteristic unbranched inflorescence called the spike, on which the floral units called spikelets are arranged in a distichous pattern. Barley produces three spikelets (one central and two lateral) per inflorescence node, and the spikelets are determinate that they form a single floret per spikelet. Whereas, wheat forms one spikelet per node, while its spikelet develops florets indefinitely without limited by a terminal floret since wheat spikelet is indeterminate (Bonnett, 1966; Kirby and Appleyard, 1984; Koppolu and Schnurbusch, 2019). Based on the fertility of the lateral spikelets, barley is commonly classified as two-rowed (both the lateral spikelets are sterile and form no grains) and six-rowed (all three spikelets are fertile and form grains). In principle, six-rowed types produce three times more grains per spike than a two-rowed spike. The major gene responsible for the row-type difference is known as *Six-rowed spike 1* (*Vrs1*) that affects the pistil development of the lateral spikelets (Komatsuda et al., 2007; Sakuma et al., 2013; Zwirek et al., 2018). Even though two-rowed lateral spikelets do not set grains, they are retained as partially developed spikelets on the two-rowed spike.

In the field of barley and hexaploid wheat (*Triticum aestivum* L.), efforts have been undertaken to unravel the impact of many traits like spike biomass, floret survival, crop growth rate, ovary size, grain number, and grain set on grain yield (Arisnabarreta and Miralles, 2006b; Arisnabarreta and Miralles, 2008a; Sadras and Slafer, 2012; Serrago et al., 2013; Alqudah and Schnurbusch, 2014; Arisnabarreta and Miralles, 2015; González-Navarro et al., 2015; Guo and Schnurbusch, 2015; Slafer et al., 2015; Gonzalez-Navarro et al., 2016; Guo et al., 2016; Garcia et al., 2019). Grain yield is majorly composed of grain number (GN) per unit area and grain weight, and it is well documented that GN is the most dominant trait influencing yield in barley and wheat (Prystupa et al., 2004; Peltonen-Sainio et al., 2007; Ugarte et al., 2007; Serrago et al., 2013; García et al., 2015; Sakuma and Schnurbusch, 2020). In these crops, the GN is determined by the number of fertile florets (wheat) or spikelets (barley) at anthesis. In general, three critical phases of the spikelet initiation and growth explains the whole process of a spikelet to grain transformation. The first phase is the floret/spikelet initiation phase that starts with the initiation of florets/spikelets and ends with the maximum number of florets/spikelets (also known as potential florets/spikelets). The end of the floret/spikelet initiation phase is known as the maximum yield potential stage (MYP). The phase following the MYP is the spike growth phase that overlaps with the stem elongation period. During this phase, the final number of florets/spikelets are determined after their abortion. The spike growth phase ends at or around the anthesis stage when the final fertile floret/spikelet number is attained. After these two phases, the fertile florets/spikelets enter the grain set phase during which the final GN is reached by filling the stored assimilates to the fertile florets/spikelets (Gallagher, 1979; Sreenivasulu and Schnurbusch, 2012).

From the available literature on wheat yield and yield components, we found two proposed models that determine the number of fertile florets or GN. The first and the most popular one is the ‘survival model’ which claims that the number of fertile florets is determined by floret survival, regulated through the assimilate allocation to the growing spike (Fischer and Stockman, 1980; Kirby, 1988; Ghiglione et al., 2008; Gonzalez et al., 2011; Ferrante et al., 2013). The other hypothesis challenged the survival model and proposed a ‘developmental model’ in which GN has a stronger correlation to the time of floret death, which is a pure developmental process. It was also shown that the onset of floret abortion is loosely associated with spike growth; instead, it is determined by the developmental stage of the most advanced floret of the spike (Bancal, 2008, 2009). Concertedly, the survival model, strongly proposes that the second phase, spikelet growth, define GN. In contrast, the developmental model supports the first spikelet initiation phase, which determines the final GN. Many studies in barley explored different yield components and their interaction with GN (del Moral et al., 1991; Prystupa et al., 2004; Ugarte et al., 2007; García et al., 2015; Kennedy et al., 2018). Also, these interactions were compared between the two- and six-rowed types (del Moral et al., 2003; Arisnabarreta and Miralles, 2006a, 2008b; Arisnabarreta and Miralles, 2008a, 2015). However, to our knowledge, there are no comprehensive studies that compared the interaction of the three phases of the spikelet initiation and growth and their interaction to the final GN in two- and six-rowed barley types. This knowledge gap must be addressed because both the row-types have a different pattern of spikelet initiation and growth and uniformity of grain size and content (Kirby and Riggs, 1978; Riggs and Kirby, 1978; Frégeau-Reid et al., 2001; Zwirek et al., 2018; Dodig et al., 2019; Kandic et al., 2019).

In this study, we explored the interaction of the critical yield components like potential spikelet number (PSN), spikelet survival (SSL), and grain set (GS) with GN from the main culm of barley row-types. Also, we analyzed the association of a few more traits (from the main culm), such as spikelet number (SN) at harvest and grain survival (GSL) with GN to understand their impact on the final GN in barley. We compared these interactions between the barley row-types, two- and six-rowed, which were grown in the greenhouse and field. Interestingly, two-rowed types performed consistently better in the field for all the traits except PSN, while in six-rowed only GN and GSL were improved. The interactions of PSN, SSL and GS with GN in both growth conditions clearly distinguished the row-types and revealed that the two-rowed GN strongly depends on PSN rather than SSL and GS. Contrastingly, in six-rowed, GN is majorly influenced by SSL than PSN and GS. The GS in six-rowed exhibited a much higher impact on GN in the greenhouse than in the field, while in two-rowed GN is less affected by GS in both the environments. Collectively, from this study, we concluded that barley row-types are distinct in their yield components interaction with GN. Specifically, in two-rowed, GN depends on the spikelet initiation trait, PSN, while in six-rowed, it is majorly influenced by the spikelet growth trait, SSL.

## Materials and Methods

### Plants and growth conditions

The study was carried out at the Leibniz Institute of Plant Genetics and Crop Plant Research, Gatersleben, Germany (51° 49′ 23″ N, 11° 17′ 13″ E, altitude 112 m) during the 2017 growing season. Twenty-seven barley accessions (Supplementary Table 1) were selected from a worldwide collection panel (Alqudah et al., 2014) based on their variation in spike developmental traits. The selected panel was used for two experiments conducted in the field (April to August 2017) and greenhouse (July to December 2017). For both the experiments, similar sowing and pre-treatment protocols were followed. Barley grains were sown on 96 well trays and grown under greenhouse conditions (photoperiod, 16h/8h, light/dark; temperature, 20° C/16° C, light/dark) for two weeks. Plants were vernalized at 4° C for four weeks and then acclimatized in the greenhouse for a week. Following the hardening process, plants were directly transplanted in the silty loam soil for the field experiment. The field experiment was a single plot per genotype, and each plot had eight rows with a 15 cm distance between the rows. Each row was 0.8 m long with five plants per row. In the greenhouse experiment (photoperiod, 16h/8h, light/dark; temperature, 20° C/16° C, light/dark), plants were potted in a 9 cm pot (9 × 9 cm, diameter × height) with two parts of autoclaved compost, two parts of ‘Rotes Substrat’ (Klasmann-Deilmann GmbH, Germany), and one part of sand. We conducted a third experiment in the field (same location mentioned above) named ‘Random culm-field’ between April 2019 to August 2019. In this experiment, the 27 genotypes were replicated in three random block model, in which a block size was 1 × 1.5 m^2^, and in every block, a genotype was sown with a density of 300 grains/m^2^. Each block had six rows spaced 20 cm between them. Standard practices for irrigation, fertilization, and control of pests and diseases were followed. The temperature, light, and other environmental conditions of the field, greenhouse, and random culm-field experiments are given in supplementary table 2.

### Phenotyping traits and measurement

All the traits reported from the field and greenhouse studies (the year 2017) were measured from the main culm of a barley plant because of its higher contribution to the final grain yield and it is less influenced by environmental perturbations and growth conditions (Cottrell et al., 1985; Elhani et al., 2007). The direct yield component traits like potential spikelet number (PSN), final spikelet number (SN), and grain number (GN) were collected from randomly selected plants at two stages, maximum yield potential (MYP) and harvest. PSN was counted at the MYP stage, while SN and GN were enumerated at the harvest. At the MYP stage, data were collected from three random plants for both the experiments, whereas at harvest, six plants were used in the field and twelve plants in the greenhouse experiment. However, in the random culm-field experiment, all the parameters were collected from three random culms of every block, which made nine replications per genotype. The MYP stage was identified by tracking the development of spikes based on the developmental scales previously been used (Waddington, 1983; Kirby and Appleyard, 1984). The stage at which the initiation of spikelet primordia stops at the apex of the main shoot spike was considered as the MYP stage.

All the initiated spikelet primordia and differentiated spikelets were counted at the MYP stage and regarded as potential spikelet number (PSN) of a spike. At MYP, more apically localized spikelet primordia ridges in the double ridge stage were multiplied by three since barley spikes form three spikelet primordia at a spike axis (rachis) node (Waddington, 1983; Kirby and Appleyard, 1984). The final SN and GN were counted from a spike after the physiological maturity of the plant. All (assimilate-) filled and unfilled spikelets were counted as final SN, while only the filled spikelets were regarded as grains. Spikelet and grain survival traits were calculated from the equations (1) and (2), respectively, and grain set from equation (3).

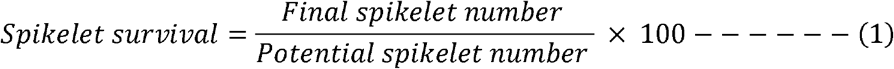

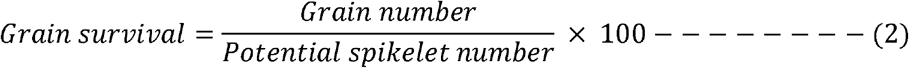

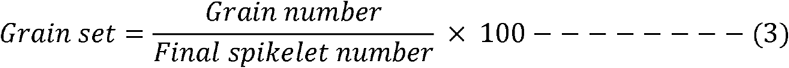

### Methods of data analysis

All the data analyzes were done in the Prism software, version 8.4.2 (GraphPad Software, LLC), except the non-linear quadratic (squared) fit. The outliers were identified by the ‘ROUT’ method described previously (Motulsky and Brown, 2006). The mean value comparison between the environments for every genotype and row-types was analyzed by the multiple t-tests and paired t-test (parametric), respectively. The false discovery rate approach of the two-stage linear step-up procedure of Benjamini, Kreier, and Yekutieli with the Q value of 5% was applied to calculate the significance of t-tests. Only for PSN and GS, a two-way ANOVA with Tukey’s multiple comparison test (alpha=5%) was conducted. The replicate values of a genotype and the row-types were analyzed individually, without assuming a consistent standard deviation. Linear regression was done using the appropriate dependent (Y values) and independent (X values) traits. The 95% confidence intervals were identified for every linear regression and plotted as confidence bands along with the ‘goodness of fit’ line. Non-linear quadratic (squared) fit for selected dependent and independent traits was done with the data analysis tool kit of Microsoft office-Excel (version 2016).

## Results

### Potential spikelet number is influenced by the combination of genetic and environmental variations

Previous studies in barley did not consider lateral spikelet primordia and spikelets of two-rowed barleys as “potential spikelets” for the determination of MYP (Arisnabarreta and Miralles, 2004, 2006b). However, after the discovery of several *Vrs1* alleles (Komatsuda et al., 2007; Sakuma et al., 2017; Casas et al., 2018) and various row-type genes that influence lateral spikelet initiation and growth (Ramsay et al., 2011; Koppolu et al., 2013; Bull et al., 2017; van Esse et al., 2017; Youssef et al., 2017; Zwirek et al., 2018), it was clearly shown that the two-rowed lateral spikelets, though sterile, also influence grain size of the central spikelets. To account for this deficiency, we included lateral spikelet primordia and spikelets of two-rowed types as potential spikelets besides the central spikelets for main culm MYP determination. Within the selected 17 two-rowed types, the genotype BCC1705 developed the lowest PSN (75±0) in the greenhouse, while BCC929 produced the highest PSN (140±1.7) in the field. Comparably, among the ten chosen six-rowed genotypes, BCC768 developed the lowest PSN in the greenhouse (77±4.5), and BCC192 formed the highest PSN (123±0) in the field (Fig.1B). Interestingly, only two genotypes from the selected six-rowed types produced PSN more than 120 (Fig. 1B), while, in two-rowed, eight genotypes showed PSN greater than or equal to 120 (Fig.1A). This indicates that in our selected panel of row-types, more two-rowed genotypes produced higher PSN than six-rowed genotypes. Additionally, the highest PSN among the row-types is also formed by a two-rowed genotype, BCC929 (Fig. 1A). Two-rowed genotypes, BCC801, HOR2828, BCC1433, and Hockett, formed significantly less PSN in the field compared to the greenhouse. However, eight of the two-rowed types, BCC1440, BCC929, PI467826, HOR18914, BCC1705, BCC1707, Metcalfe, and Bowman, developed significantly higher PSN in the field. The rest of the two-rowed types showed similar (i.e., statistically non-significant) PSN in the two environments (Fig.1A). Correspondingly, in the six-rowed genotypes, BCC149 developed significantly less PSN in the field, while BCC719 and Morex produced more PSN in the field than the greenhouse. The remaining six-rowed genotypes had a similar number of potential spikelets in both environments (Fig.1B). Comparing the mean phenotypic value of the two environments showed that both the row-types have similar mean PSN (Fig. 1C), and they are not significantly different between environments. However, the PSN comparison of every genotype displayed more than half of the genotypes (12 two-rowed and three six-rowed) had a significant difference between the growth conditions (Fig. 1A & B). This result suggested that in barley, PSN is combinatorially influenced by genetic and environmental variations.

**Figure 1.**
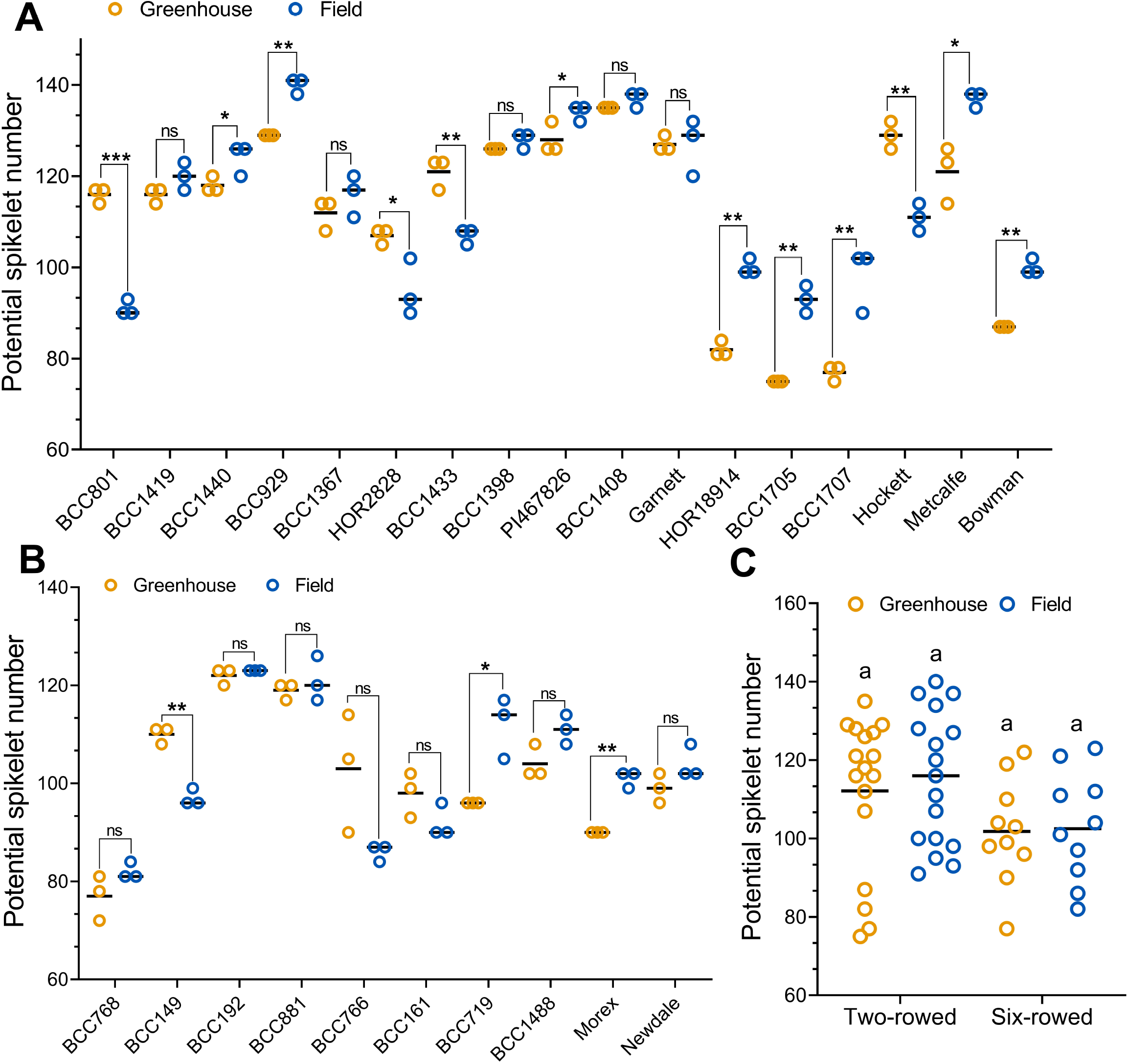
Potential spikelet number variation of row-types in the greenhouse and field. A comparison of the main culm potential spikelet number (PSN) between greenhouse and field conditions of two-rowed (A) and six-rowed barleys (B) is shown. A) Two-rowed genotypes BCC801, HOR2828, BCC1433, and Hockett showed a reduction, whereas BCC1440, BCC929, PI467826, HOR18914, BCC1705, BCC1707, Metcalfe, and Bowman exhibited an increase in PSN in the field. B) In six-rowed, only BCC149 developed a reduced PSN in the field compared to the greenhouse. Contrarily, genotypes BCC719 and Morex formed more PSN in the field. C) A comparison of the mean values of the PSN between greenhouse and field of two- and six-rowed genotypes is shown. PSN is not significantly different between row-types and the environments. In A & B, each genotype was represented by three replicates both in the greenhouse and field. In C, the mean values of 17 two-rowed and ten six-rowed genotypes were used. Data in A & B were analyzed by *multiple Student’s t-tests* with false discovery analysis of *Benjamini, Kreier, and Yekutieli* with the *q* value of 5% and the data in C with *Two-way ANOVA* with *Tukey’s multiple comparison test (alpha=5%)*; Letter ‘a’ in panel C denotes that the mean values are not significantly different at *q*>0.05. *, *q*≤0.05; **, *q*≤0.01; ***, *q*≤0.001; ns, *q>0.05*.

### Survival traits of two-rowed barleys are highly influenced by the environment

The survival traits measure the ability of a plant to produce spikelets (SSL) or grains (GSL). Unlike PSN, SSL and GSL cannot be calculated for every replicate of a genotype because counting the PSN at the MYP stage is a destructive approach. Therefore, it is not possible to relate the spikelet or GN of a replicate to its PSN, and so only the mean values are used for those two traits (Fig 2 & 3). In two-rowed, all genotypes either maintained similar ability or showed a higher level of main culm SSL or GSL in the field (Fig. 2A & 3A). Only one accession, the six-rowed type Morex, reduced its survival traits in the field compared to the greenhouse (Fig. 2B & Fig. 3B). Two-rowed types BCC1440 and BCC1408, had a similar level of SSL, and the rest of the other genotypes had higher SSL in the field (Fig. 2A). Additionally, all the two-rowed genotypes (including BCC1440 and BCC1408) displayed a greater GSL in the field compared to the greenhouse (Fig. 3A). In the six-rowed types, except for Morex, all other genotypes either had similar (BCC768 and BCC766) SSL and GSL in both the environments or had an increasing trend in the field compared to the greenhouse (Fig. 2B & Fig. 3B). In both row-types, the lowest SSL and GSL were formed in the greenhouse and the highest was produced in the field, indicating that field conditions are more amenable for both row-types to exhibit higher survival (spikelet and grain) abilities. The two-rowed genotype BCC801 had the lowest SSL (16.5 %) and GSL (12.6 %), while BCC1433 possessed the highest SSL (28.3 %) and GSL (24.9 %) (Fig. 2A & 3A). Among the six-rowed genotypes, BCC149 had the lowest SSL (45.6 %), while BCC881 had the lowest GSL (26.1 %). The six-rowed genotypes BCC1488 showed the highest SSL (75.2 %), while Newdale had the highest GSL (63 %) (Fig. 2B & Fig. 3B). We did not compare the absolute values of the survival traits between the row-types because two-rowed types, in general, abort two-thirds of their spikelets anyhow (apart from the apical abortion) due to the influence of *Vrs1* gene function (Komatsuda et al., 2007; Sakuma et al., 2013; Zwirek et al., 2018). The average ability of SSL and GSL comparison of the two environments showed that the two-rowed types exhibited higher SSL and GSL in the field, while the six-rowed types only had greater GSL in the field. Additionally, two-rowed types consistently displayed a higher significance (*q*<0.001) for both traits in the field (Fig. 2C & 3C), which indicated that the two-rowed barleys’ survival traits (spikelet and grain) are more highly influenced by the environment.

**Figure 2.**
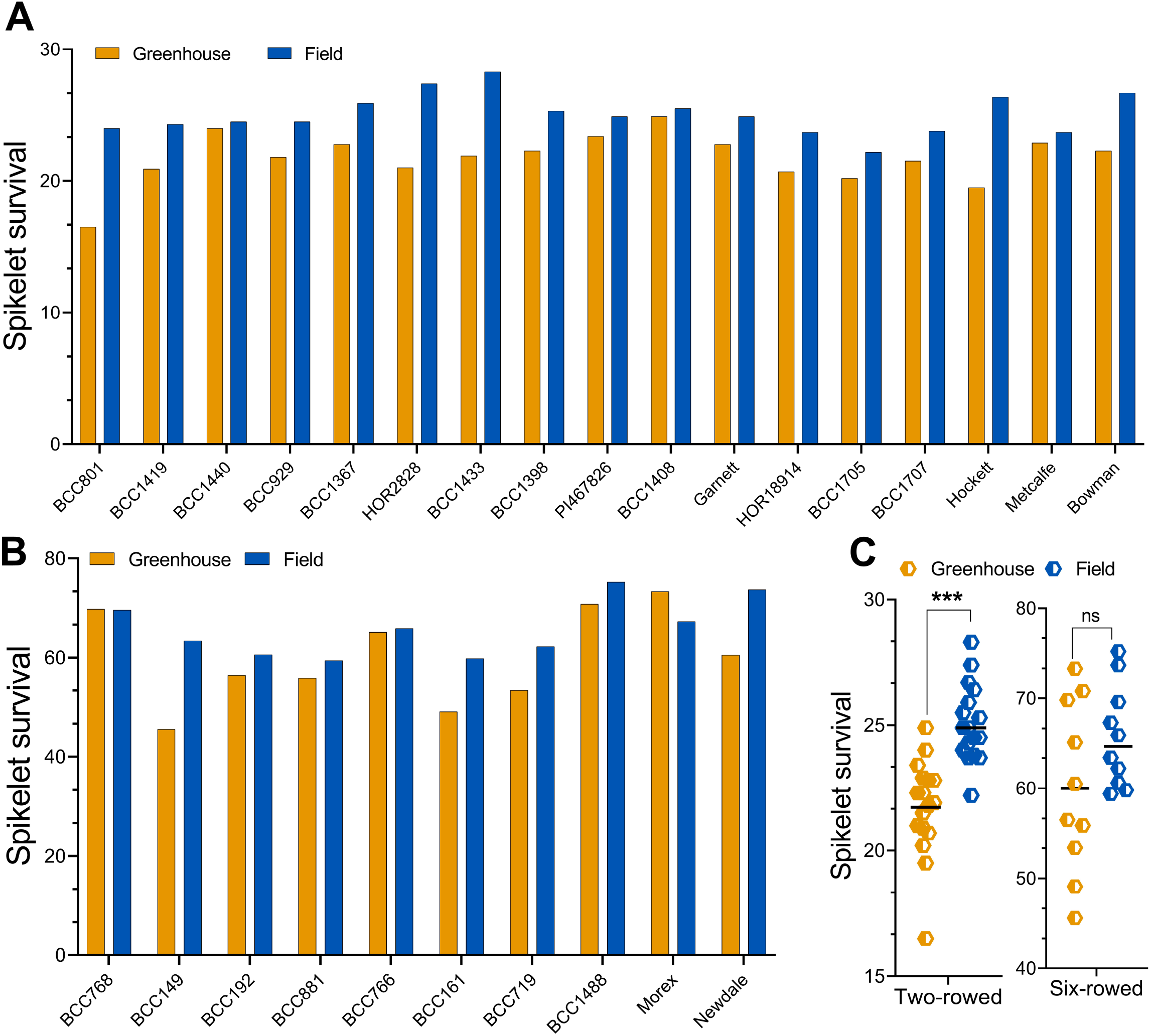
Spikelet survival variation of row-types in the greenhouse and field. A comparison of the mean main culm spikelet survival (SSL) between the greenhouse and field conditions of two-rowed (A) and six-rowed barleys (B) is shown. A) In two-rowed, except BCC1440 and BCC1408, all other genotypes exhibited a higher trend of SSL in the field. B) In six-rowed, genotypes BCC149, BCC192, BCC881, BCC161, BCC719, BCC1488, and Newdale showed a trend of increased SSL in the field compared to the greenhouse while Morex had a decreased trend in the field. C) A comparison of the mean values of SSL between the greenhouse and field of two- and six-rowed genotypes is shown. Two-rowed showed an increased variation of SSL in the field, while six-rowed exhibited similar variation between the environments. In A & B, each genotype was represented by the mean value from 4 to 12 replicates in the greenhouse and six replicates in the field. In C, the mean values of 17 two-rowed and 10 six-rowed genotypes were used. In C, the data were analyzed by *paired Student’s t-test (parametric)* with false discovery analysis of *Benjamini, Kreier, and Yekutieli* with the *q* value of 5%; ***, *q*≤0.001; ns, *q>0.05*.

**Figure 3.**
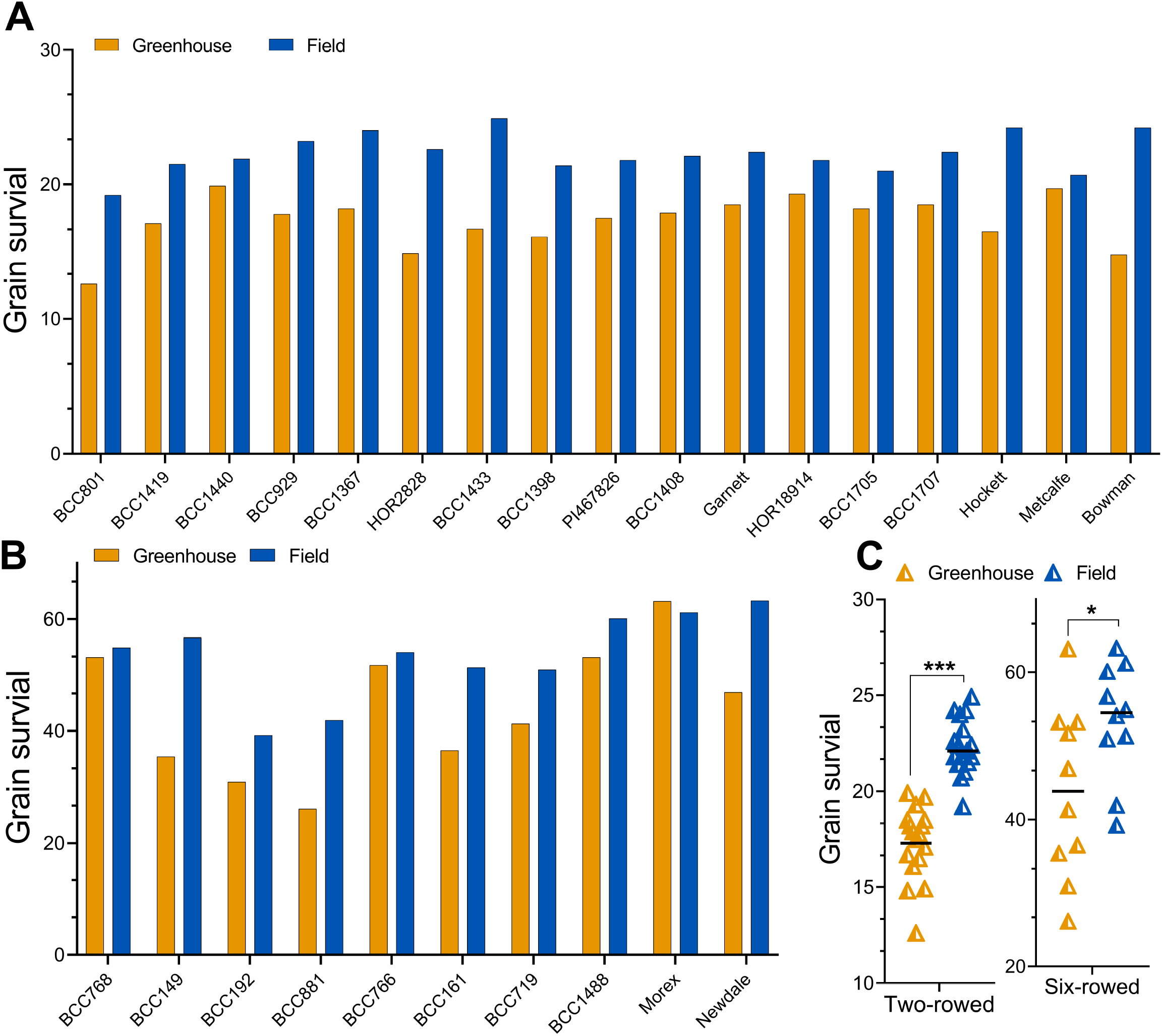
Grain survival variation of row-types in the greenhouse and field. A comparison of the mean main culm grain survival (GSL) between the greenhouse and field conditions of two-rowed (A) and six-rowed barleys (B) is shown. A) In two-rowed, all genotypes exhibited a higher trend of grain survival in the field. B) In six-rowed, except Morex that showed a reduced SSL and GSL in the field, the rest of other genotypes either had similar (BCC768 and BCC766) GSL in both the environments or had an increasing trend of GSL in the field compared to the greenhouse. C) A comparison of the mean values of GSL between greenhouse and field of two- and six-rowed genotypes is shown. Both row-types showed increased GSL in the field, while two-rowed genotypes showed a more significant increase than six-rowed. In A & B, each genotype was represented by the mean value from 4 to 12 replicates in the greenhouse and six replicates in the field. In C, the mean values of 17 two- and ten six-rowed genotypes were used. In C, the data were analyzed by *paired Student’s t-test (parametric)* with false discovery analysis of *Benjamini, Kreier, and Yekutieli* with the *q* value of 5%; ***, *q*≤0.001; *, *q*≤*0.05*.

### Two-rowed barleys exhibited a higher grain set than six-rowed types

Grain set (GS) estimates the potential of a plant to transform the final fertile spikelets into grains successfully. Within our panel, all genotypes either had a similar or higher main culm GS under field conditions (Fig. 4A-C), which reinforced the fact that in the field, our panel of barley row-types performed better than in the greenhouse. The two-rowed genotypes, BCC929, BCC1367, PI467826, BCC1408, BCC1705, and Bowman, as well as the six-rowed types, BCC149, BCC192, BCC881, and BCC161, had a significant increase of GS in the field compared to the greenhouse (Fig. 4A & B). The remaining six- and two-rowed genotypes maintained a similar, i.e., non-significant, GS between the environments. Also, for GS, like SSL and GSL, we observed the lowest GS of both row-types in the greenhouse, while the greatest was in the field. Within two-rowed types, Bowman showed the lowest (66.43±9.07 %) and BCC929, BCC1705, and BCC1707, the highest (~94%) GS (Fig. 4A). Similarly, in the six-rowed types, BCC881 had the lowest (46.91±10.47 %), while Morex and BCC149 had the highest (~90%) GS (Fig. 4B). In two-rowed, many genotypes, like BCC1440, BCC929, BCC1367, Garnett, HOR18914, BCC1705, BCC1707, Hockett, and Bowman had greater than or equal to 90% GS in the field compared to two six-rowed genotypes (BCC149 and Morex) (Fig. 4A & B). This suggested that GS of the two-rowed types exhibited a more significant response in the field, which is also reflected in the mean GS comparison between row-types and environments (Fig. 4C). Here, two-rowed types exhibited significantly (*q*=0.005) higher GS ability in the field compared to the greenhouse. Moreover, the two-rowed average GS ability was higher than in the six-rowed types in the greenhouse (*q*<0.0001) and field (*q*=0.04) (Fig. 4C), revealing that two-rowed barleys exhibit the best GS ability after all.

**Figure 4.**
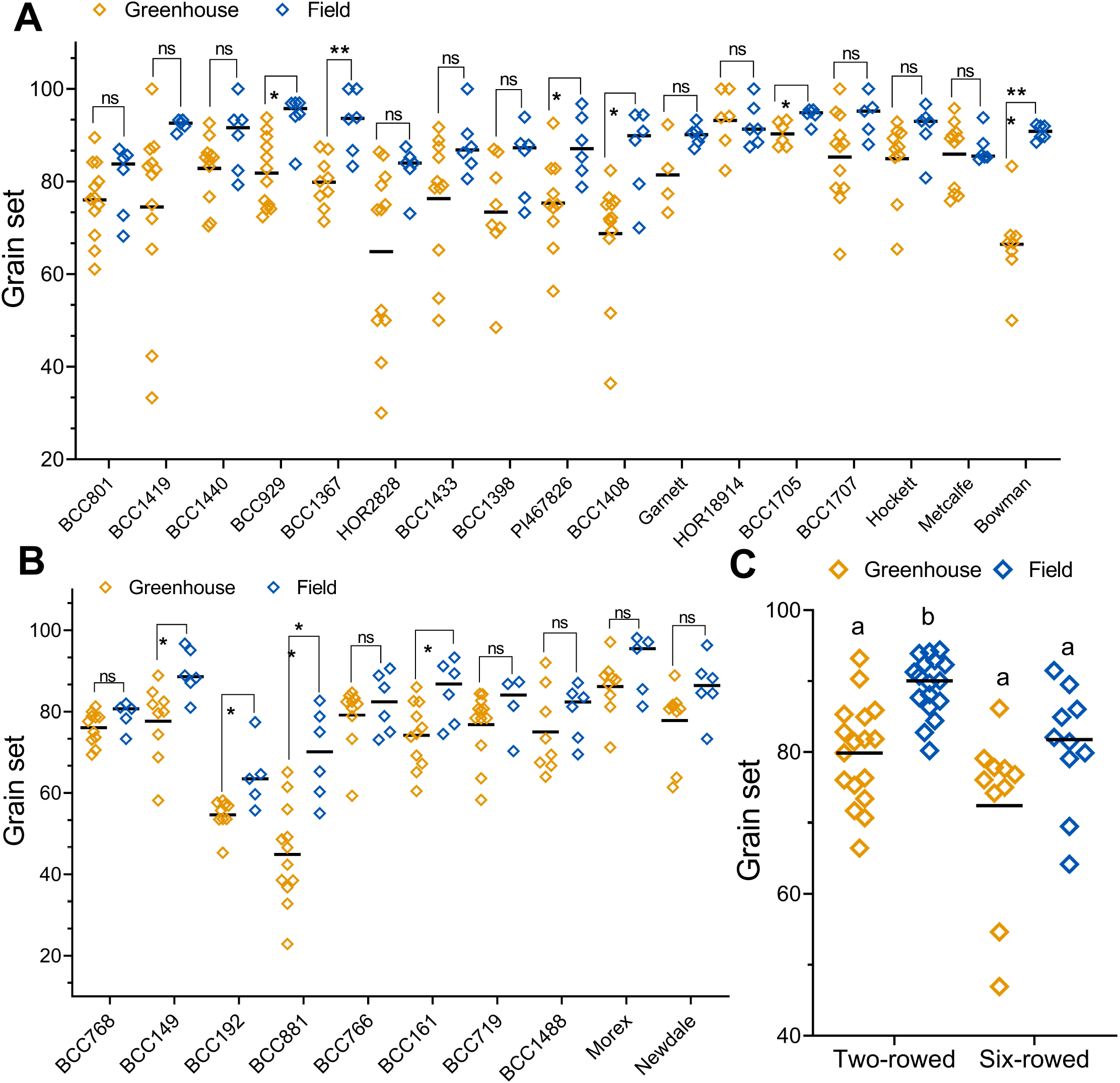
Grain set variation of row-types in the greenhouse and field. A comparison of the main culm grain set (GS) between greenhouse and field conditions of two-rowed (A) and six-rowed barleys (B) is shown. A) In two-rowed, genotypes BCC929, BCC1367, PI467826, BCC1408, BCC1705, and Bowman showed higher GS in the field. B) In six-rowed, genotypes BCC149, BCC192, BCC881, and BCC161 had significantly higher GS in the field compared to the greenhouse. C) A comparison of the mean values of GS between the greenhouse and field of the two- and six-rowed genotypes is shown. Two-rowed showed an increased variation of GS in the field, while six-rowed exhibited similar variation between the environments. In A & B, each genotype was represented by 4 to 12 replicates in the greenhouse and six replicates in the field. In C, the mean values of 17 two- and ten six-rowed genotypes were used. Data in A & B were analyzed by *multiple Student’s t-tests* with false discovery analysis of *Benjamini, Kreier, and Yekutieli* with the *q* value of 5% and the data in C with *Two-way ANOVA* with *Tukey’s multiple comparison test (alpha=5%)*; Letter ‘b’ in panel C shows that the two-rowed field GS is significantly different than two-rowed greenhouse (*q*≤0.001), six-rowed greenhouse (*q*≤0.001), and six-rowed field (*q*≤0.05). However, the letter ‘a’ denotes that the mean values are not significantly different at *q*>0.05. *, *q*≤0.05; **, *q*≤0.01; ***, *q*≤0.001; ns, *q>0.05*.

### Spikelet and grain number are differentially regulated by the environment in barley row-types

In our two-rowed panel, except for BCC1408 that had similar SN of the main culm between the environments, all others developed a significantly higher SN in the field (Fig. 5A). Interestingly, all two-rowed genotypes, including BCC1408, formed significantly more grains (main culm) in the field compared to the greenhouse (Fig. 6A). Conversely, the six-rowed genotype BCC766 developed significantly fewer spikelets and grains in the field (Fig. 5B & 6B). The remaining six-rowed types either maintained similar or an increased SN (Fig. 5B) and GN (Fig. 6B) in the field compared to the greenhouse. Here also, for both row-types, we found that the lowest SN and GN were in the greenhouse and the highest in the field. In two-rowed, the lowest SN (19.2±1.02) was formed by BCC801 and the highest (35±3.1) by BCC1408 (Fig. 5A). Similarly, for GN, the lowest (12.9±2) was produced by Bowman and the highest (32.5±1.8) by BCC929 (Fig. 6A). Comparably, the six-rowed genotypes BCC161 had the lowest (48.1±5.7), and BCC1488 had the highest SN (83.5±3.6) (Fig. 5B). The six-rowed genotype BCC881 formed the lowest GN (31±6.5) and the highest by BCC1488 (66.7±6.1). The average phenotypic value of the SN and GN comparison of the two environments showed that only the two-rowed types developed significantly more SN (Fig. 5C) and GN (Fig. 6C) in the field. In contrast, six-rowed types produced only higher GN in the field. Similar to the findings in SSL, GSL, and GS, we observed that two-rowed types had a consistently higher increase of SN and GN in the field (Fig. 5C & 6C). From this result, it becomes evident that the environment (growth conditions) affects GN of both row-types, but SN only in two-rowed types.

**Figure 5.**
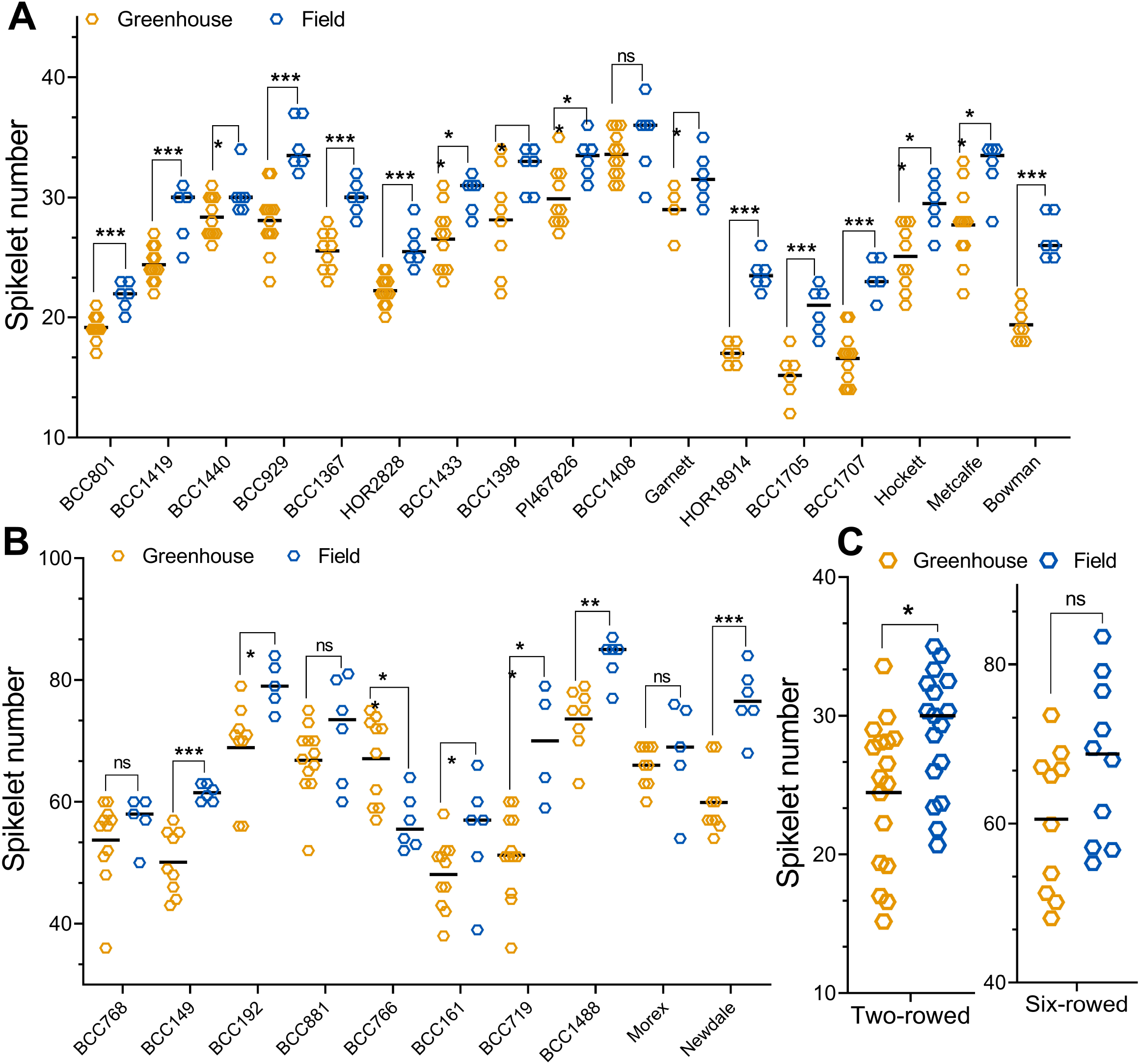
Final spikelet number variation of row-types in the greenhouse and field. A comparison of the main culm spikelet number (SN) between greenhouse and field conditions of two-rowed (A) and six-rowed barleys (B) is shown. A) Except for the two-rowed type BCC1408 that have similar SN between the environments, all others were developed a significantly higher SN in the field. B) In six-rowed, BCC766 developed significantly less SN in the field. At the same time, the rest of the other genotypes either maintained similar (BCC768, BCC881, & Morex) or produced significantly higher (BCC149, BCC192, BCC161, BCC719, BCC1488, and Newdale) SN in the field. C) A comparison of the mean values of the SN between the greenhouse and field of two- and six-rowed genotypes is shown. Two-rowed showed significantly higher variation in the field while six-rowed had similar variation in both growth conditions. In A & B, each genotype was represented by 4 to 12 replicates in the greenhouse and six replicates in the field. In C, the mean values of 17 two- and ten six-rowed genotypes were used. Data in A &B were analyzed by *multiple Student’s t-tests* and the data in C with *paired Student’s t-test* (*parametric*) with false discovery analysis of *Benjamini, Kreier, and Yekutieli* with the *q* value of 5%; *, *q*≤0.05; **, *q*≤0.01; ***, *q*≤0.001; ns, *q>0.05*.

**Figure 6.**
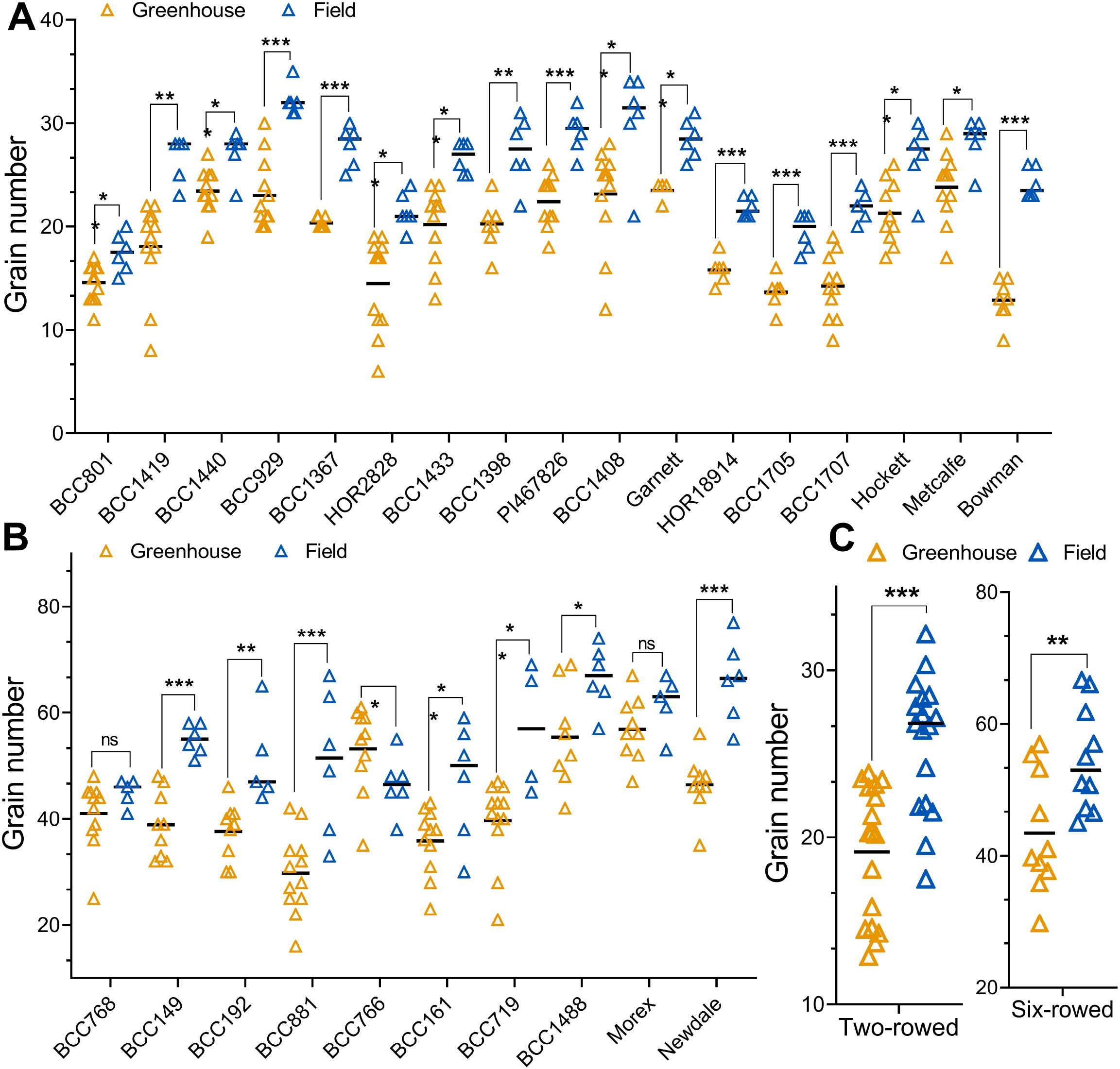
Grain number variation of row-types in the greenhouse and field. A comparison of the main culm grain number (GN) between greenhouse and field conditions of two-rowed (A) and six-rowed barleys (B) is shown. A) All two-rowed genotypes formed a higher GN significantly in the field compared to the greenhouse. B) The six-rowed genotype BCC766 produced significantly lower GN in the field while the rest of the other genotypes either maintained similar (BCC768, Morex) or higher GN in the field. C) A comparison of the mean values of GN between greenhouse and field of two- and six-rowed genotypes is shown. Both row-types developed significantly more grains in the field. In A & B, each genotype was represented by 4 to 12 replicates in the greenhouse and six replicates in the field. In C, the mean values of 17 two- and ten six-rowed genotypes were used. Data in A &B were analyzed by *multiple Student’s t-tests* and the data in C with *paired Student’s t-test* (*parametric*) with false discovery analysis of *Benjamini, Kreier, and Yekutieli* with the *q* value of 5%; *, *q*≤0.05; **, *q*≤0.01; ***, *q*≤0.001; ns, *q>0.05*.

### Two and six-rowed types have distinct interactions of yield components for main culm grain number determination

To reveal the processes influencing main culm GN determination in barley, we assessed the interaction between main culm PSN, SSL, GS, and GN of two- and six-rowed barleys in both environments. In the greenhouse, the GN of the two-rowed was strongly dependent on PSN (R^2^=0.68, *P*<0.0001) and moderately on SSL (R^2^=0.35, *P*=0.012) (Fig. 7A & B). Interestingly, in the field, the interaction between GN and PSN became stronger (R^2^=0.86, *P*<0.0001), while the GN and SSL turned weaker (R^2^=0.06, *P*=0.356) (Fig. 8A & B). In two-rowed, GN seems independent from the effect of GS in both environments (Fig. 7C & 8C). Contrastingly under greenhouse conditions, the six-rowed GN was predominantly determined by SSL (R^2^=0.56, *P*=0.012) and GS (R^2^=0.49, *P*=0.025) (Fig. 7E & F). These interactions were significantly reduced in the field. Nevertheless, the SSL (R^2^=0.39, *P*=0.052) ended up as a major yield component, (among other traits) deciding the six-rowed GN (Fig. 8E & F). In both the environments (growth conditions), the six-rowed GN was independent of the influence on PSN (Fig. 7D & 8D). From these results, it is clear that the row-types follow unique yield component interactions for regulating their main culm GN.

**Figure 7.**
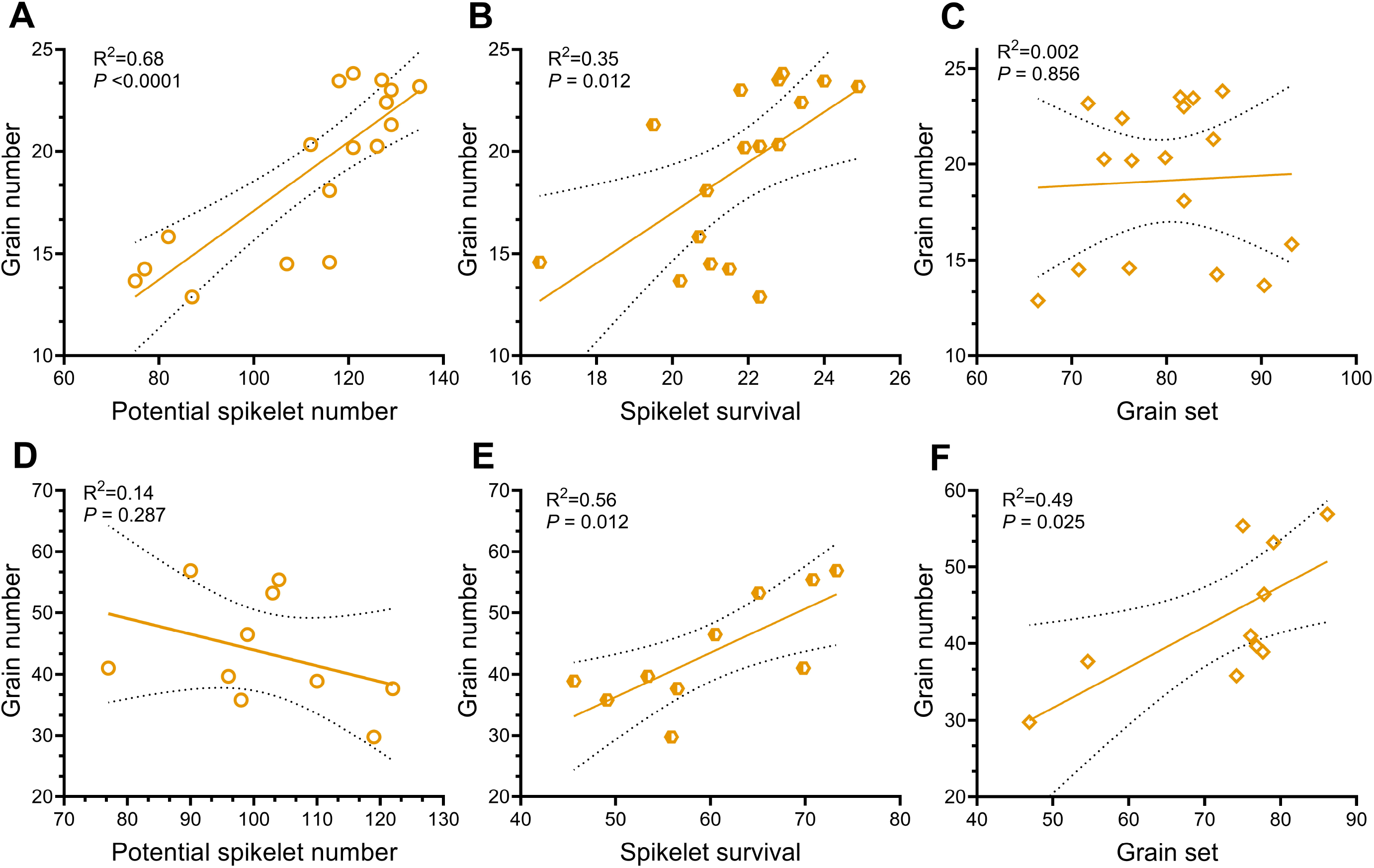
Interactions of potential spikelet number, spikelet survival, and grain set with grain number in the greenhouse. The interactions of potential spikelet number (PSN) with grain number (GN), spikelet survival (SSL) with GN, and grain set (GS) with GN of two- (A, B, & C) and six-rowed (D, E, & F) are shown respectively. In two-rowed, GN was predominantly determined by PSN, while in six-rowed by SSL and GS. Linear regression analysis was done by considering GN as dependent and PSN or SSL or GS as independent traits for both row-types. 95% confidence intervals were plotted as confidence bands.

**Figure 8.**
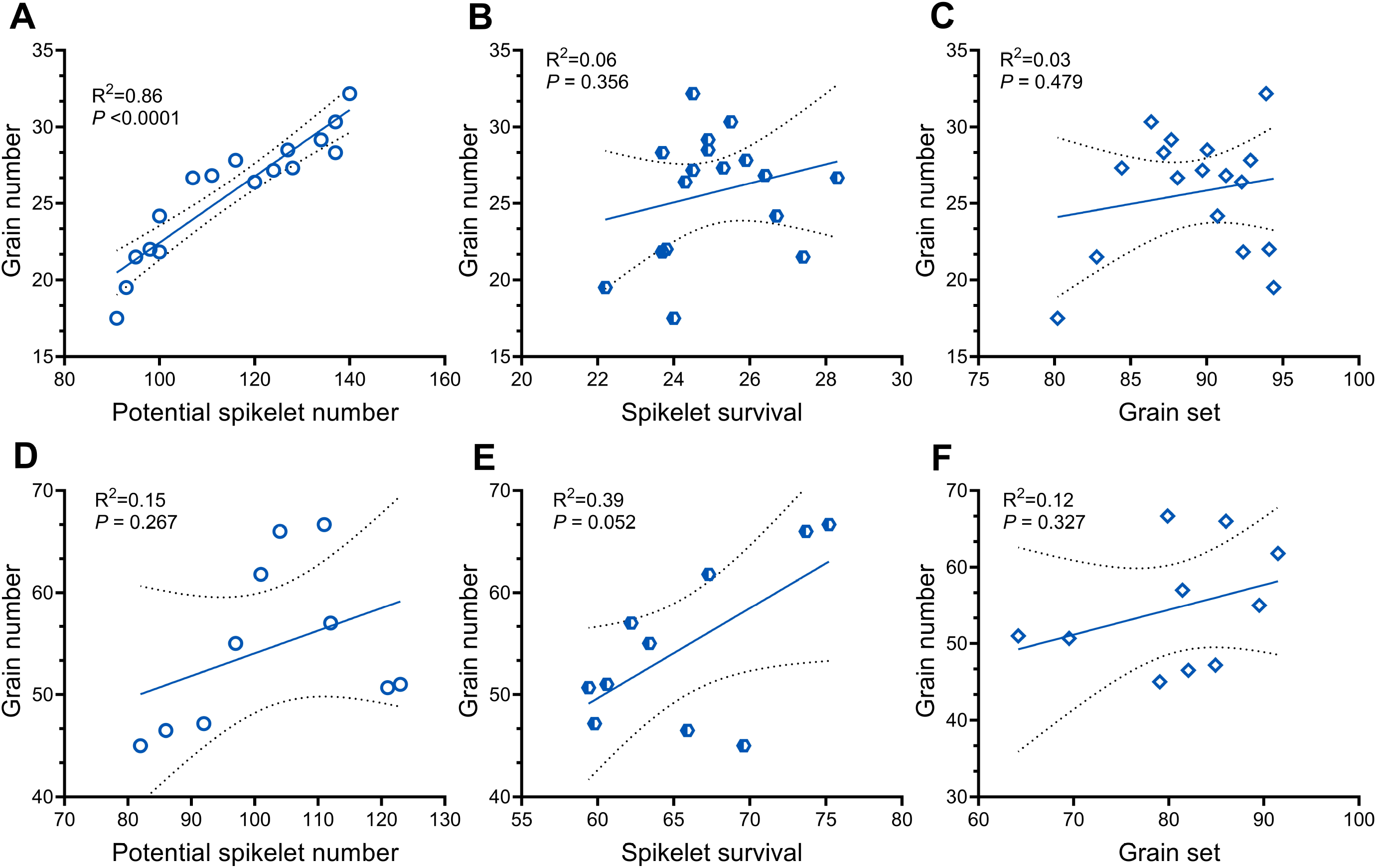
Interactions of potential spikelet number, spikelet survival, and grain set with grain number in the field. The interactions of potential spikelet number (PSN) with grain number (GN), spikelet survival (SSL) with GN, and grain set (GS) with GN of two- (A, B, & C) and six-rowed (D, E, & F) are shown respectively. In two-rowed, GN was predominantly determined by PSN, while in six-rowed by SSL. Linear regression analysis was done by considering GN as dependent and PSN or SSL or GS as independent traits for both row-types. A 95% confidence interval was plotted as confidence bands.

## Discussion

The study was aimed to understand better the interactions between main culm spikelet initiation and growth processes in relation to the main culm GN determination in two- and six-rowed barley. We compared these interactions in two different environments, controlled greenhouse, and field, to assess the stability of their associations. Intriguingly, the PSN was found to be influenced both by the genetical and environmental variations in barley. This observation partly supports the hypothesis that in barley, PSN is a pure developmental factor that is highly controlled by a genetic component, as previously claimed by another study (Arisnabarreta and Miralles, 2006a).

### Two-rowed barley follows the developmental model for grain number determination

In two-rowed, GN was predominantly determined by PSN and to a lesser extent by SSL and GS in both the environments (Fig. 7A & 8A), suggesting that the interaction between PSN and GN is highly stable. Additionally, the above result implied that two-rowed genotypes developed higher PSN at the end of the spikelet initiation phase, produced more grains at harvest compared to the genotypes with fewer PSN, irrespective of their SSL and GS. A similar result was reported in an earlier study (Digel et al., 2015) with a two-rowed barley cv. Scarlett, in which the reduction of the early reproductive phase with a dominant *Ppd-H1* allele significantly diminished the potential spikelets and the number of final floret/spikelet primordia. We also found that the interactions of GN with SN and SN with PSN were robust for two-rowed in both growth conditions (Fig. S1A-D). These interactions reiterated that in two-rowed, GN is mainly determined by the variation in PSN through SN. Field studies on two- and six-rowed barley predicted that the critical period for yield determination could be earlier in two-rowed than six-rowed types (Arisnabarreta and Miralles, 2006b; Arisnabarreta and Miralles, 2008a). Our study also supports this view because we found that the two-rowed GN is predominantly determined by PSN, which is the product of the spikelet initiation phase. Conversely, six-rowed GN had a strong association with SSL, which is a process that happens during the spikelet growth phase, following the spikelet initiation phase. A moderate linear relationship between SSL and GN only in the greenhouse suggested that the two-rowed spikelets competed either with main culm growth or whole plant tiller development during stem elongation, which was not the case in the field (Ghiglione et al., 2008; Gonzalez et al., 2011; Ferrante et al., 2013; Guo and Schnurbusch, 2015; Guo et al., 2017; Guo et al., 2018a; Guo et al., 2018b; Guo et al., 2018c; Perez-Gianmarco et al., 2019). The disconnection of GS with GN in both the environments (Fig. 7C & 8C) suggested that assimilates stored during the pre-anthesis phase were sufficient for grain-filling of two-rowed fertile spikelets. Thus, there is less competition between the spikelets of the two-rowed main culm spike during the GS phase. Collectively, our results supported that the GN determination in two-rowed accessions largely depends on the developmental trait PSN; however, we also found some exceptions in the field. Two-rowed SSL (R^2^=0.30, *P*=0.08) and GS (R^2^=0.35, *P*=0.05) showed a moderate non-linear quadratic (squared) relationship with GN in the field (Fig. S2 A&B), suggesting that the field conditions affected specific two-rowed genotypes that showed either a positive or negative association with GN. Further analysis of these genotypes may reveal more intriguing interactions and influences of SSL and GS to GN. We also verified the interactions in another field study, in which we collected data from random culms instead of from the main culm. Intriguingly, in addition to the strong influence (R^2^=0.77, *P*<0.0001) of PSN on GN, we found that GS had significant (R^2^=0.76, *P*<0.0001) interaction to GN (Fig. S3 A-C). This suggested that in two-rowed different culms might have additional influences in determining GN. Thus, our study exemplified that two-rowed PSN strongly influences GN and that the GN of the two-rowed barley types may follow the developmental model (Bancal, 2008, 2009).

### Six-rowed grain number follows the survival model

In general, the products of the spikelet initiation phase, otherwise known as PSN, enters into the spikelet growth phase during which some potential spikelets or/and spikelet primordia are aborted, most likely and at least partly, due to the competition with the developing main culm and tillers. The end products of the spikelet growth phase are the fertile spikelets (SN) that are filled in the GS phase with assimilates preserved during pre-anthesis. There is also another kind of abortion (known as grain abortion) that happens during the GS phase due to the limited availability and/or allocation of assimilates to the fertile spikelets (Gallagher, 1979). Contrasting to the two-rowed types, six-rowed GN showed a moderate interaction with SSL; but was lowered for PSN both in the greenhouse (Fig. 7D & E) and field (Fig. 8D & E). Also, six-rowed GN exhibited an association with GS only in the greenhouse (Fig 7F). Similar interactions were reported from a field study with the near-isogenic lines of two- and six-rowed barley, demonstrating that the critical period for determining the yield components of six-rowed rather occurred during the survival and part of GS phase (Arisnabarreta and Miralles, 2008a). Also, many studies from wheat claimed that the floret survival period and the competition between florets and other vegetative organs determine final fertile florets (Kirby, 1988; Ghiglione et al., 2008; Gonzalez et al., 2011; Ferrante et al., 2013; Guo and Schnurbusch, 2015; Guo et al., 2017; Guo et al., 2018a; Guo et al., 2018b; Guo et al., 2018c; Perez-Gianmarco et al., 2019). Thus, our results reiterated that the GN of the six-rowed mostly depends on the survival processes irrespective of the growth conditions. It also implied that six-rowed spikelets are highly competitive with other initiation and/or growth events that happen during the spikelet survival phase. The strong interaction of six-rowed SSL with GN indicated that genotypes with high spikelet survival (low spikelet abortion) produce more grains at harvest. The above result is in line with a previous study (Arisnabarreta and Miralles, 2008a), predicting that the critical period for setting grains in six-rowed barley falls within 30 days before heading. A further enhancement of GN potential in six-rowed might be possible if one modulates the survival process (Arisnabarreta and Miralles, 2008a, 2010, 2015), which, however, requires further studies dissecting the genetic modulators of spikelet survival in six-rowed barley. The inconsistent association of GS to GN between the experiments (Fig. 7F & 8F) suggested that the plasticity of GS might be under the environmental influence, which supports the previous outcome observed in a six-rowed field study (Voltas et al., 1997). We only observed an exception in the field conditions, where six-rowed GN showed a non-linear quadratic (squared) (R^2^=0.73, *P*=0.011) relationship with PSN (Fig. S2C). This indicates that similar to the two-rowed SSL and GS, the field condition impacted certain six-rowed types that showed positive or negative interaction of GN with PSN. We also compared the observations to another field study, in which the random culms of six-rowed types had weak interactions (R^2^=0.39, *P*=0.055) between PSN and GN, and the influence of SSL on GN was lost (R^2^=0.14, *P*=0.0291). However, GS showed a strong interaction (R^2^=0.77, *P*=0.0008) with GN (Fig. S3 D-F). These results suggested that in six-rowed, culms developed in different time windows might have various interactions with GN, as observed in two-rowed barleys. In summary, our results supported that main culm GN determination in six-rowed barley genotypes depends on the SSL; and thus, it rather follows the survival model (Kirby, 1988; Ghiglione et al., 2008; Gonzalez et al., 2011; Ferrante et al., 2013; Perez-Gianmarco et al., 2019).

## Conclusion

Our study revealed the intrinsic interactions of main culm yield components such as PSN, SSL, and GS with GN of two- and six-rowed barleys in two different growth conditions. We found that GN in two-rowed accessions is majorly determined during the early spikelet initiation phase because it has a strong interaction with PSN in both environments. The spikelets of two-rowed barleys may have moderate or less competition with the other growing parts during the survival phase due to the inconsistent and weak association of SSL with GN. This effect can be partly explained by the two-thirds of the two-rowed spikelets that do not set any grains, i.e., sterile lateral spikelets. Conversely, in six-rowed, the lateral spikelets form grains, so the GN is mainly determined by the competition of its spikelets within the spike and/or other vegetative/structural organs during the survival phase. A similar hypothesis on the influence of lateral spikelets on two- and six-rowed barley’s was predicted earlier (Cottrell and Dale, 1984; Arisnabarreta and Miralles, 2004). Thus, from this study, we therefore, propose that the barley row-types follow two distinct patterns of GN determination, where two-rowed barleys follow a developmental model, and six-rowed genotypes comply with the survival model.

## Supplementary data

Figure S1. Interactions of two-rowed potential spikelet number, final spikelet number, and grain number in greenhouse and field.

Figure S2. Interactions of potential spikelet number, spikelet survival, grain set, and grain number in greenhouse and field.

Figure S3: Interactions of random culms’ potential spikelet number, spikelet survival, grain set, and grain number in the field.

Table S1: Details of the 27 accessions used in this study.

Table S2: Growth conditions of the field and greenhouse experiments.

## Acknowledgments

We thank Angelika Pueschel, Corinna Trautewig, Mechthild Puerschel, Nandhakumar Shanmugaraj, and Roop Kamal for help with sampling and harvesting. The study received financial support from the European Research Council (ERC), grant agreement 681686 “LUSH SPIKE,” ERC-2015-CoG; HEISENBERG Program of the German Research Foundation (DFG), grant no. SCHN 768/15-1 and the IPK core budget.

## Author contributions

T.S. and V.T. conceived the idea for this experiment; V.T., and R.K., conducted the experiments; V.T. analyzed the data and wrote the manuscript; T.S. edited and reviewed the manuscript together with R.K.; T.S. supervised the study.

## Supplementary figure legends

Figure S1. **Interactions of two-rowed potential spikelet number, final spikelet number, and grain number in greenhouse and field.** The interaction of the main culm final spikelet number (SN) with grain number (GN) of two-rowed barleys in the greenhouse (A) and field (C) is shown. Similarly, the interaction of potential spikelet number (PSN) with final SN of two-rowed barleys in the greenhouse (B) and field (D) is shown. Linear regression analysis was done by considering either GN (A & C) or SN (B & D) as dependent and SN (A & C) or PSN (B & D) as independent traits for two-rowed types. 95% confidence intervals were plotted as confidence bands.

Figure S2. **Interactions of potential spikelet number, spikelet survival, grain set, and grain number in greenhouse and field.** The non-linear interaction of main culm spikelet survival (SSL) with grain number (GN) and main culm grain set (GS) with GN of two-rowed barleys in field conditions shown in A & B respectively. In C, the non-linear interaction of six-rowed main culm potential spikelet number (PSN) with GN in the field condition is shown. In A, B & C, we found a non-linear quadratic (squared) relationship with respected dependent and independent traits. The analysis was done by considering GN as dependent and SSL (A) or GS (B) or PSN (C) as independent traits.

Figure S3: **Interactions of random culms’ potential spikelet number, spikelet survival, grain set, and grain number in the field.** The interactions of potential spikelet number (PSN) with grain number (GN), spikelet survival (SSL) with GN, and grain set (GS) with GN of two- (A, B, & C) and six-rowed (D, E, & F) are shown respectively. A 95% confidence interval was plotted as confidence bands.

