## Supplementary figures and images for "Strategies of grain number determination differentiate barley row-types"

### Figure-S1

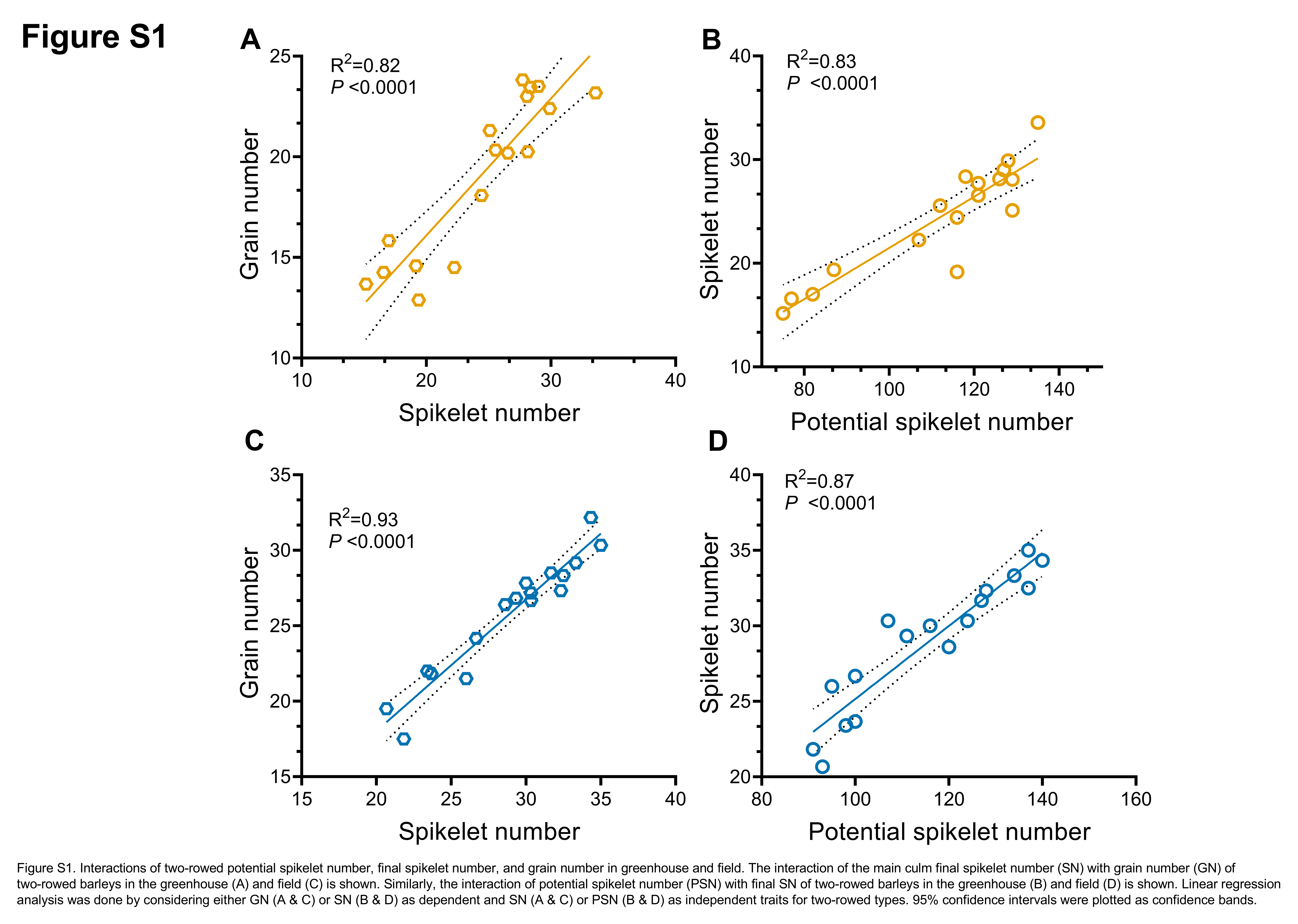

### Figure-S2

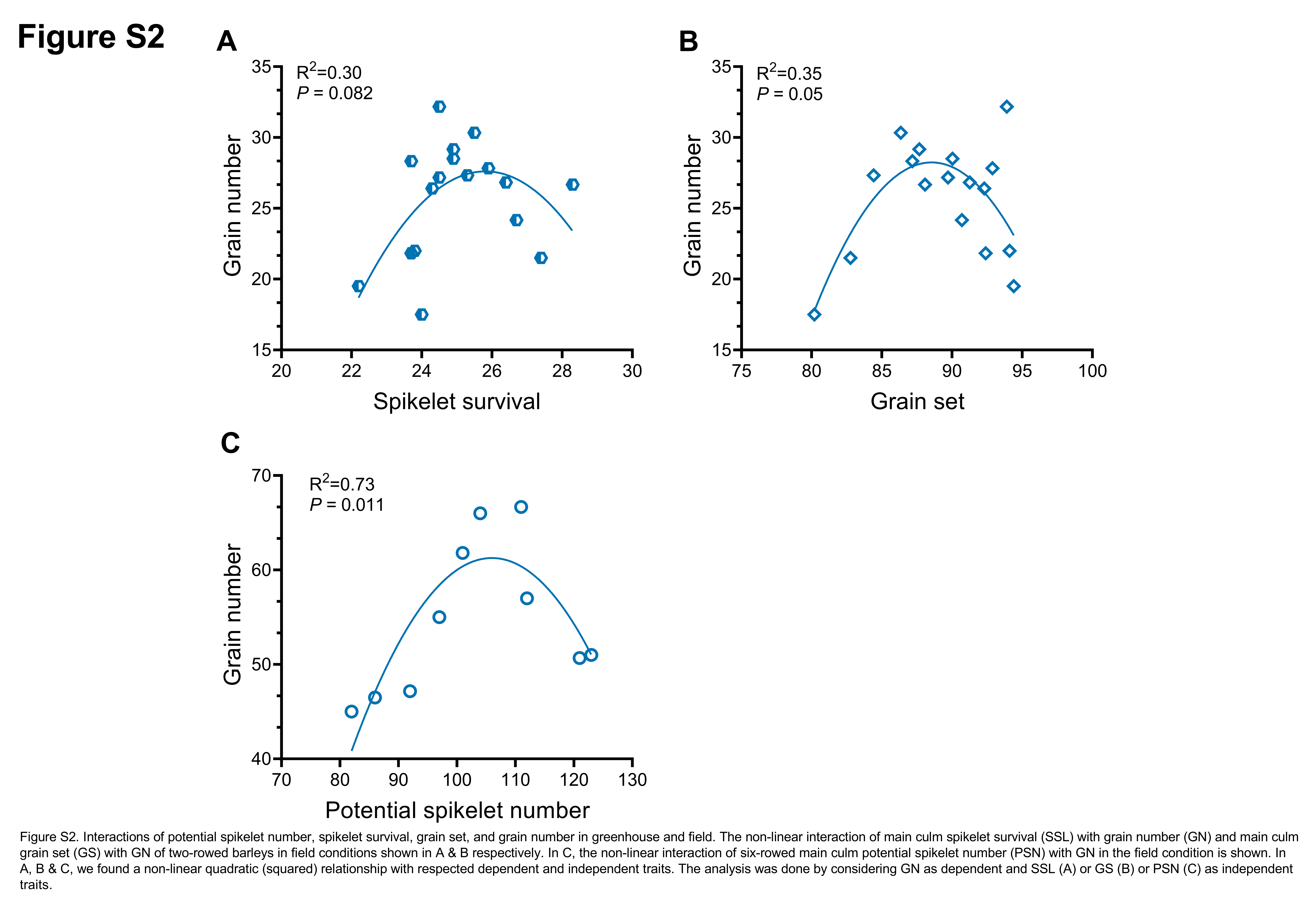

### Figure-S3

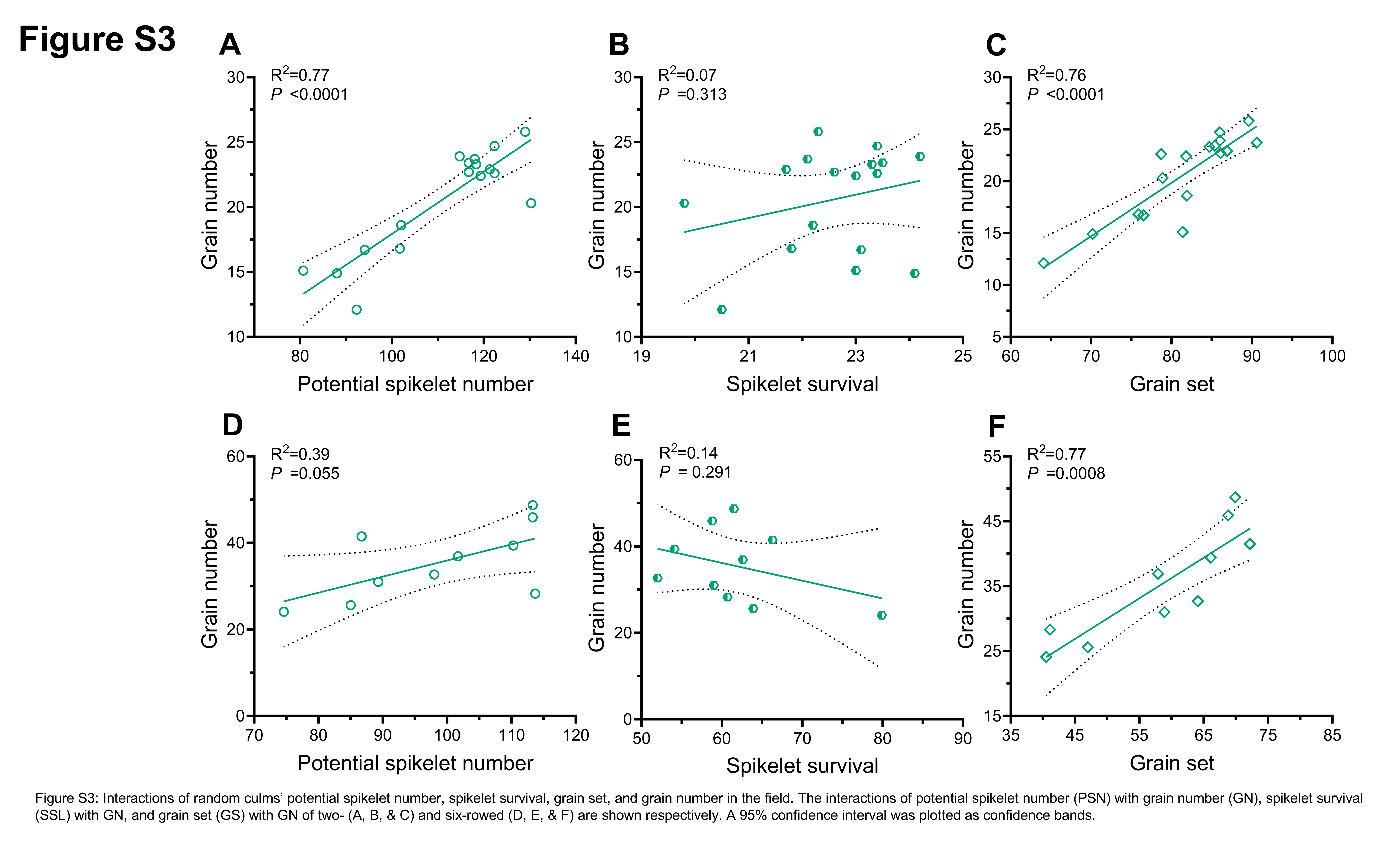
